## Supplementary figures and images for "Blood pressure variability compromises vascular function in middle-aged mice"

### Suppl Fig 1

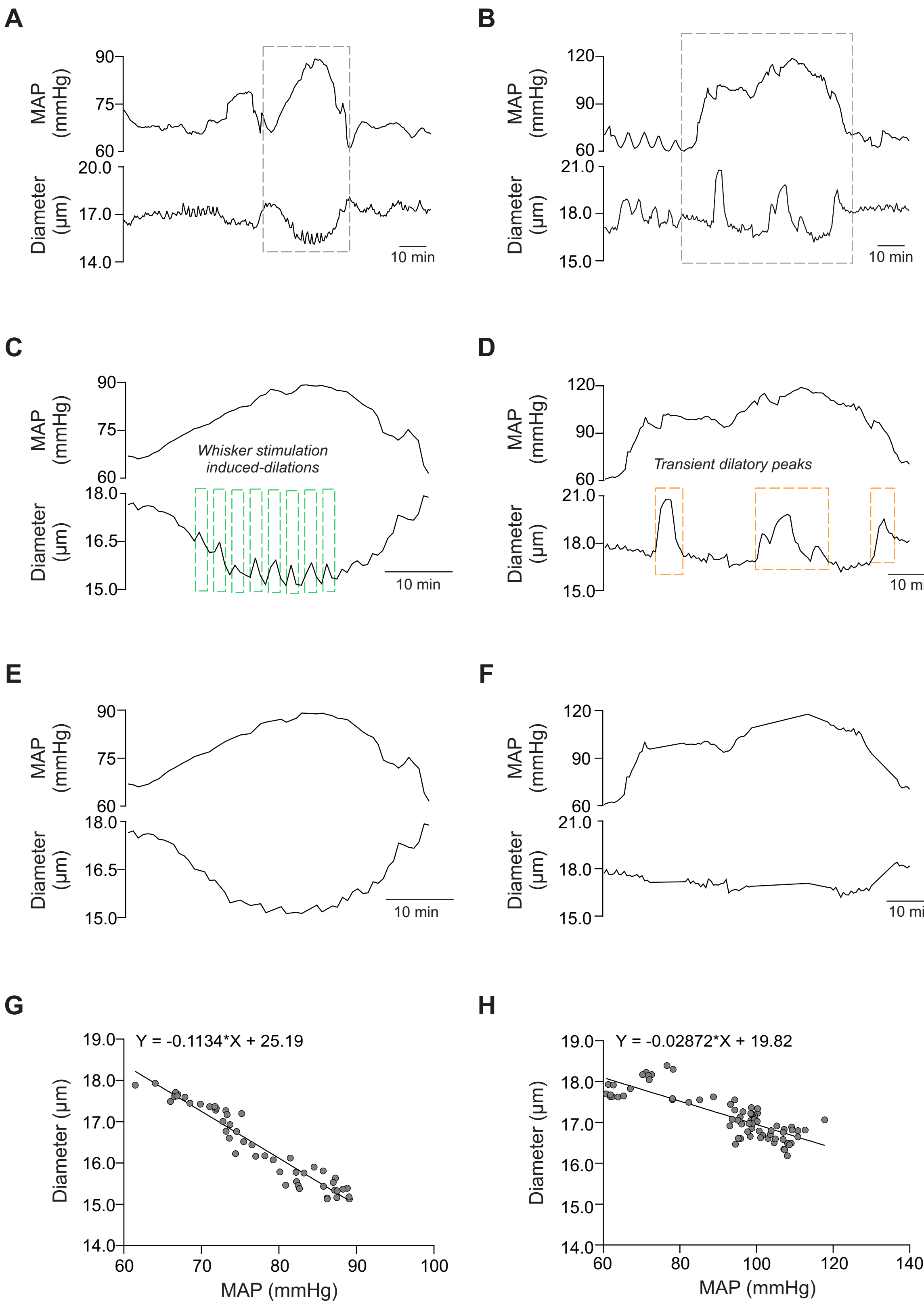

### Suppl Fig 2

Suppl. Fig 2

**A** *24-hour Averaged BP at Baseline*

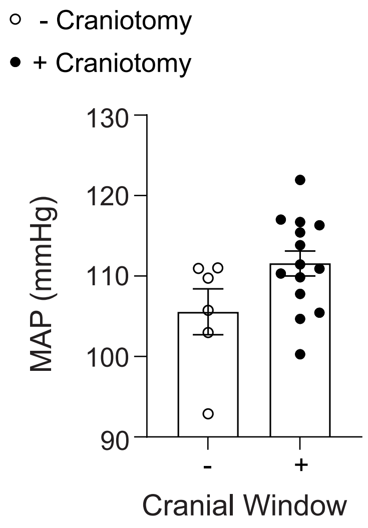


**B** *24-hour Averaged BP*

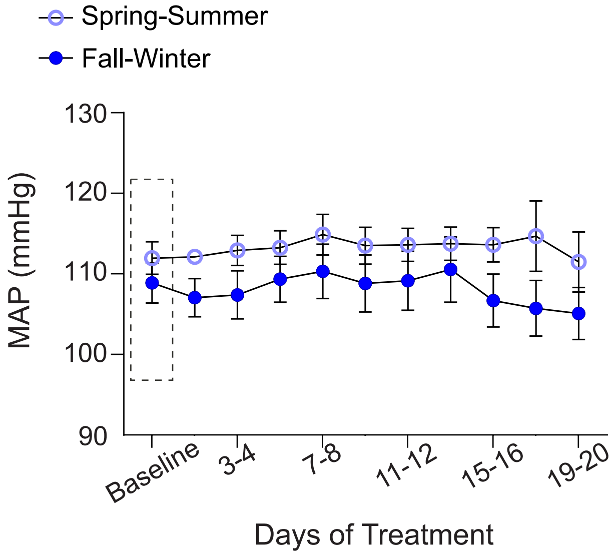

### Suppl Fig 3

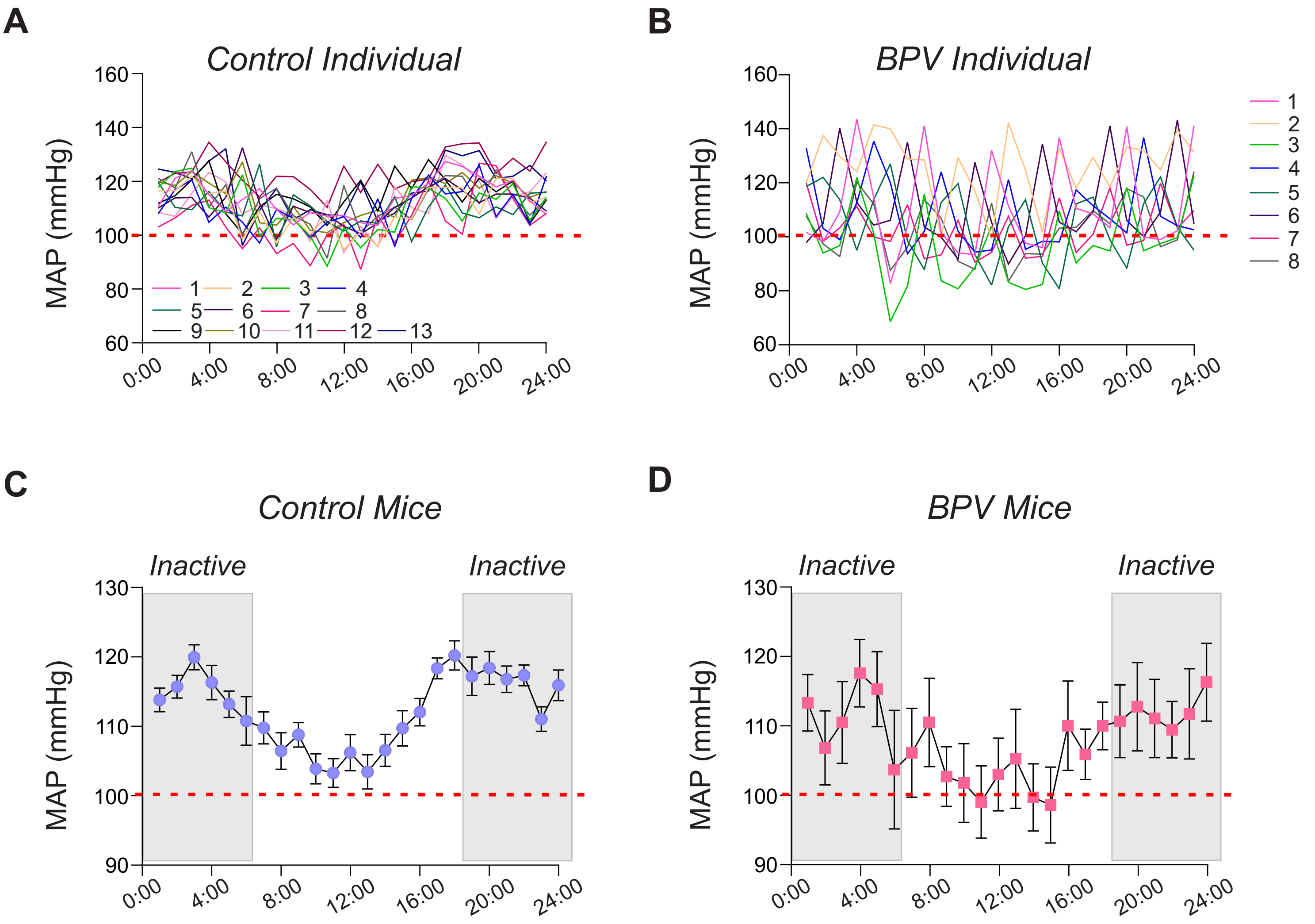

### Suppl Fig 4

#### Suppl. Fig 4

*Infusion-Evoked (Control Group)*

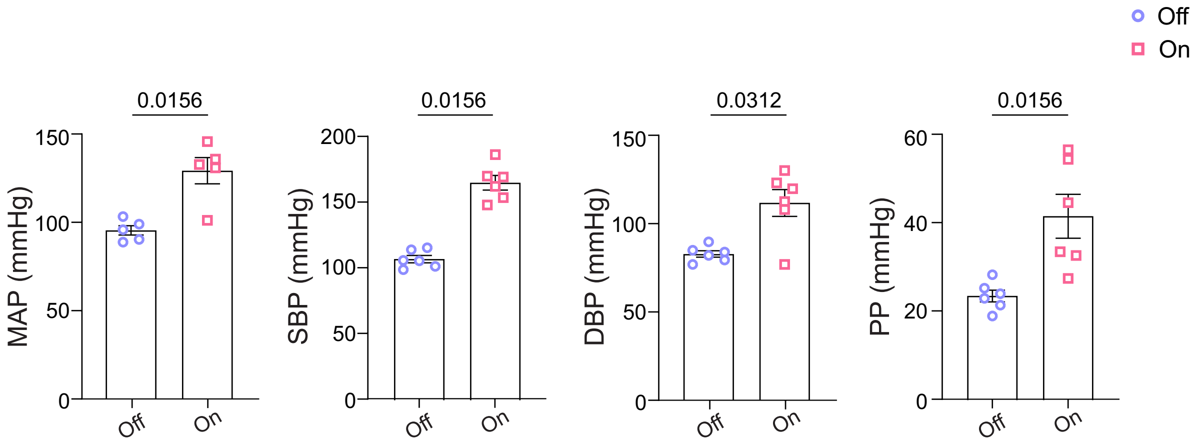

### Suppl Fig 5

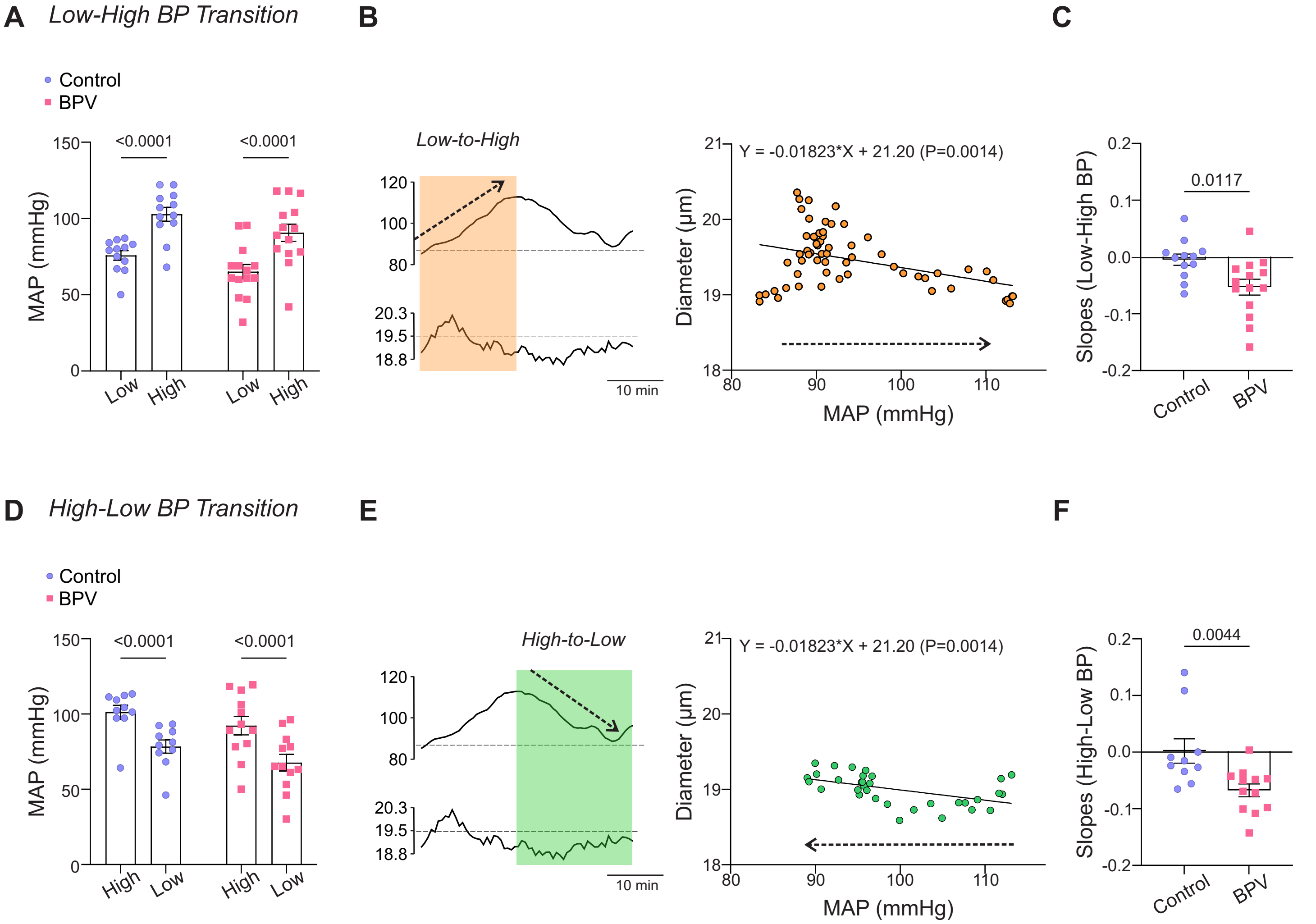
