## Supplementary material for "Blood pressure variability compromises vascular function in middle-aged mice": Suppl Table 1

**A**

### Active Period CV mmHg ± SEM

| Days | MAP | | SBP | | DBP | | PP | |
|---|---|---|---|---|---|---|---|---|
| | Control | BPV | Control | BPV | Control | BPV | Control | BPV |
| *Baseline* | *0.05 ± 0.00* | *0.05 ± 0.00* | *0.05 ± 0.00* | *0.05 ± 0.00* | *0.06 ± 0.00* | *0.05 ± 0.00* | *0.08 ± 0.01* | *0.07 ± 0.01* |
| 1-2 | 0.06 ± 0.01 | 0.09 ± 0.01 * | 0.06 ± 0.01 | 0.12 ± 0.01*# | 0.06 ± 0.01 | 0.09 ± 0.01 | 0.09 ± 0.01 | 0.22 ± 0.02 *# |
| 3-4 | 0.05 ± 0.00 | 0.11 ± 0.01 *# | 0.06 ± 0.00 | 0.13 ± 0.01*# | 0.06 ± 0.00 | 0.11 ± 0.01*# | 0.09 ± 0.01 | 0.20 ± 0.02 *# |
| 5-6 | 0.05 ± 0.00 | 0.12 ± 0.01 *# | 0.05 ± 0.00 | 0.13 ± 0.01*# | 0.05 ± 0.00 | 0.12 ± 0.01*# | 0.08 ± 0.01 | 0.20 ± 0.02 *# |
| 7-8 | 0.05 ± 0.01 | 0.12 ± 0.01 *# | 0.05 ± 0.01 | 0.14 ± 0.02*# | 0.05 ± 0.01 | 0.12 ± 0.01*# | 0.08 ± 0.01 | 0.22 ± 0.03 *# |
| 9-10 | 0.05 ± 0.01 | 0.13 ± 0.01 *# | 0.05 ± 0.01 | 0.15 ± 0.01*# | 0.05 ± 0.01 | 0.14 ± 0.01*# | 0.08 ± 0.01 | 0.21 ± 0.03 *# |
| 11-12 | 0.05 ± 0.01 | 0.13 ± 0.01 *# | 0.05 ± 0.01 | 0.15 ± 0.01*# | 0.05 ± 0.01 | 0.12 ± 0.01*# | 0.09 ± 0.01 | 0.22 ± 0.03 *# |
| 13-14 | 0.05 ± 0.01 | 0.13 ± 0.01 *# | 0.05 ± 0.01 | 0.15 ± 0.01*# | 0.05 ± 0.01 | 0.13 ± 0.01*# | 0.09 ± 0.01 | 0.24 ± 0.03 *# |
| 15-16 | 0.04 ± 0.01 | 0.14 ± 0.01 *# | 0.04 ± 0.01 | 0.16 ± 0.01*# | 0.04 ± 0.01 | 0.13 ± 0.01*# | 0.09 ± 0.01 | 0.26 ± 0.03 *# |
| 17-18 | 0.05 ± 0.00 | 0.14 ± 0.01 *# | 0.05 ± 0.00 | 0.16 ± 0.01*# | 0.05 ± 0.00 | 0.14 ± 0.01*# | 0.10 ± 0.01 | 0.26 ± 0.04 * |
| 19-20 | 0.06 ± 0.01 | 0.13 ± 0.01 *# | 0.06 ± 0.01 | 0.15 ± 0.01*# | 0.05 ± 0.01 | 0.12 ± 0.01*# | 0.12 ± 0.02 | 0.25 ± 0.02 *# |

**B**

### Inactive Period CV mmHg ± SEM

| Days | MAP | | SBP | | DBP | | PP | |
|---|---|---|---|---|---|---|---|---|
| | Control | BPV | Control | BPV | Control | BPV | Control | BPV |
| *Baseline* | *0.06 ± 0.00* | *0.06 ± 0.01* | *0.06 ± 0.00* | *0.06 ± 0.01* | *0.07 ± 0.00* | *0.06 ± 0.01* | *0.11 ± 0.01* | *0.07 ± 0.01* |
| 1-2 | 0.08 ± 0.01 | 0.10 ± 0.01* | 0.08 ± 0.01 | 0.12 ± 0.01* | 0.08 ± 0.01 | 0.10 ± 0.01* | 0.12 ± 0.03 | 0.21 ± 0.03* |
| 3-4 | 0.07 ± 0.00 | 0.13 ± 0.01*# | 0.07 ± 0.01 | 0.15 ± 0.01*# | 0.07 ± 0.01 | 0.14 ± 0.01*# | 0.12 ± 0.02 | 0.21 ± 0.02*# |
| 5-6 | 0.07 ± 0.01 | 0.14 ± 0.01*# | 0.07 ± 0.01 | 0.15 ± 0.01*# | 0.07 ± 0.01 | 0.14 ± 0.02*# | 0.12 ± 0.04 | 0.19 ± 0.02* |
| 7-8 | 0.07 ± 0.01 | 0.14 ± 0.01*# | 0.08 ± 0.01 | 0.16 ± 0.01*# | 0.07 ± 0.01 | 0.15 ± 0.01*# | 0.11 ± 0.04 | 0.23 ± 0.04* |
| 9-10 | 0.07 ± 0.01 | 0.16 ± 0.01*# | 0.07 ± 0.01 | 0.18 ± 0.02*# | 0.07 ± 0.01 | 0.17 ± 0.01*# | 0.10 ± 0.04 | 0.22 ± 0.03* |
| 11-12 | 0.07 ± 0.01 | 0.16 ± 0.01*# | 0.07 ± 0.01 | 0.18 ± 0.01*# | 0.08 ± 0.01 | 0.16 ± 0.01*# | 0.12 ± 0.03 | 0.24 ± 0.04* |
| 13-14 | 0.08 ± 0.02 | 0.16 ± 0.01*# | 0.08 ± 0.01 | 0.18 ± 0.01*# | 0.08 ± 0.01 | 0.15 ± 0.02*# | 0.12 ± 0.03 | 0.25 ± 0.04* |
| 15-16 | 0.07 ± 0.01 | 0.17 ± 0.01*# | 0.07 ± 0.01 | 0.19 ± 0.01*# | 0.07 ± 0.01 | 0.17 ± 0.01*# | 0.13 ± 0.03 | 0.30 ± 0.05* |
| 17-18 | 0.08 ± 0.01 | 0.17 ± 0.00*# | 0.08 ± 0.01 | 0.19 ± 0.01*# | 0.08 ± 0.01 | 0.18 ± 0.01*# | 0.14 ± 0.03 | 0.27 ± 0.04* |
| 19-20 | 0.07 ± 0.01 | 0.16 ± 0.01*# | 0.08 ± 0.01 | 0.18 ± 0.01*# | 0.08 ± 0.01 | 0.15 ± 0.01*# | 0.14 ± 0.04 | 0.28 ± 0.04* |
